# Phased epigenomics and methylation inheritance in a historical *Vitis vinifera* hybrid

**DOI:** 10.1101/2025.05.27.656431

**Authors:** Noé Cochetel, Amanda Vondras, Rosa Figueroa-Balderas, Joel Liou, Paul Peluso, Dario Cantu

## Abstract

Epigenetic modifications, such as DNA methylation, regulate transcription and influence key biological traits. While many efforts were made to understand their stability in annual crops, their long-term persistence in clonally propagated plants remains poorly understood. Grapevine (*Vitis vinifera*) provides a unique model, with cultivars vegetatively propagated for centuries. Here, we assembled the phased genomes of Cabernet Sauvignon and its parental lineages, Cabernet Franc and Sauvignon Blanc, using HiFi long-reads and a gene map tenfold denser than existing maps. Using three clones per cultivar, we quantifyied methylation with very consistent short- and long-read sequencing and ensured both varietal representativeness and assessment of clonal variability. We leveraged the parent-progeny sequence graph to highlight allele-specific methylation and conserved transcriptomic patterns for genes and small RNA. Such a format was essential to integrate multi-omics data and revealed that despite less clonal conservation than genetic polymorphisms, methylation marks were remarkably inherited. By further demonstrating the linear-reference limitations, we determined the correct representation of genetic variants by the sequence graph is crucial for the accurate allelic quantification of the methylome. These findings reveal the remarkable stability of epigenetic marks in a model propagated by asexual reproduction. Using a phased sequence graph, we introduce a scalable framework that accounts for genomic variation, accurately quantifies allele-specific methylation, and supports multi-omics integration such as our evaluation of the transcriptional impact of epigenetic inheritance. This approach has broad implications for perennial crops, where epigenetic variation could influence traits relevant to breeding, adaptation, and long-term agricultural sustainability.

## Introduction

Genetic and epigenetic marks shape genome function and regulate key biological processes, influencing traits such as development, stress responses, and adaptation [1]. Among these, DNA methylation plays a critical role in regulating gene and transposable elements (TEs) expression, impacting how organisms respond to their environment and maintain cellular identity across generations [2,3]. In perennial plants, epigenetic modifications are particularly important as they affect phenotypes and plant adaptability over long lifespans [4]. Many fruit trees and other horticultural crops are propagated vegetatively to preserve desirable traits and ensure genetic and phenotypic consistency across generations. Without sexual reproduction to reshuffle genetic material, clones largely retain the genetic landscape of their progenitors, raising questions about the long-term stability of epigenetic information and how it evolves independently of genetic changes [5].

The accurate quantification of an individual’s methylome strongly depends on the concomitant availability of its genome sequence and the proper resolution of its ploidy at the haplotype level. Using a single reference genome is known to have a profound impact when characterizing genetic polymorphisms, and methylation analysis is the most affected [6]. By sequencing and phasing individual genomes, we can overcome the inherent biases of classic single-reference analyses, enabling a more accurate representation of polymorphisms, structural variants, gene content, and epigenetic marks. Beyond individual genome assemblies, pangenome graphs have emerged as a powerful approach to overcome the limitations of linear references, integrating multiple genomes into a unified framework that captures structural variation and epigenetic diversity [7]. The graphical format not only overcomes the single-reference bias but also integrates phasing information by storing phased haplotypes.

Grapevine (*Vitis vinifera*) has a long history of domestication which involved transitions of reproductive system from the wild species, obligate outcrossers, to the monoecious cultivated grapes (*Vitis vinifera* spp. *vinifera*). Despite having the potential to self-pollinate, cultivation of domesticated grapes primarily relies on hybridization and clonal propagation to avoid inbreeding depression. The vegetative propagation of cultivars contributes to the fixation of desirable phenotypes by conserving highly heterozygous mutations beneficial to grape cultivation [8]. While genomic diversity has been characterized among grape clones, to what extent the epigenetic landscape is inherited during cultivar propagation remains largely unknown [9]. One of the most famous cultivars for winemaking, Cabernet Sauvignon, is derived from a cross between Cabernet Franc and Sauvignon Blanc. The hybridization event is suggested to have occurred spontaneously around the 17^th^ century [10]. Such a model presents an outstanding opportunity to assess the persistence of parental genomic and epigenetic signatures in a well-established, clonally propagated lineage.

In this study, we assembled and annotated the genomes of Cabernet Sauvignon, Cabernet Franc, and Sauvignon Blanc to investigate to what extent methylation marks are conserved in a vegetatively propagated model. The varietal representativeness of the data relied on the usage of three independent clones as biological replicates and the methylomes were supported by both short- and long-read sequencing. By accurately phasing the genomes using a high-density gene-based map, we compared Cabernet Sauvignon to its parental lineage haplotypes, enabling us to track allele-specific methylation, gene expression, and miRNA-mRNA networks. By applying a sequence graph approach to this trio, we not only provided new insights into the long-term stability of epigenetic inheritance in a clonally propagated hybrid but also overcame the bias inherent in single-reference approaches. Our findings not only advance the study of grapevine epigenomics but also offer a broader integrative framework via sequence graphs for investigating epigenetic conservation in genomic trios formed through hybrid crossings.

## Results

### Phased genome assembly and methylation quantification for the Cabernet Sauvignon parent-progeny trio

The Cabernet Sauvignon genome was assembled by combining HiFi sequencing data, a high-density gene map, and parental lineage information (Table 1). The gene map was developed using several *Vitis vinifera* genomes selected for the high contiguity of their assembly. About 20,000 genes were designated as highly conserved markers which represents a tenfold increase in density compared to the classic rhampseq markers [11] commonly used for scaffolding with HaploSync [12]. In addition to longer reads and a richer scaffolding map, sequencing data from Cabernet Franc (CF) and Sauvignon Blanc (SB) were also used to improve the phasing and assign a parent of origin to each haplotype (i.e. hapCF and hapSB instead of hap1/2). Moreover, new assemblies for CF and SB were generated using HiFi reads with the high-density gene map resulting in significantly improved statistics when compared with previous assembly versions of the same cultivars [13,14] (Table 1).

**Table 1.**
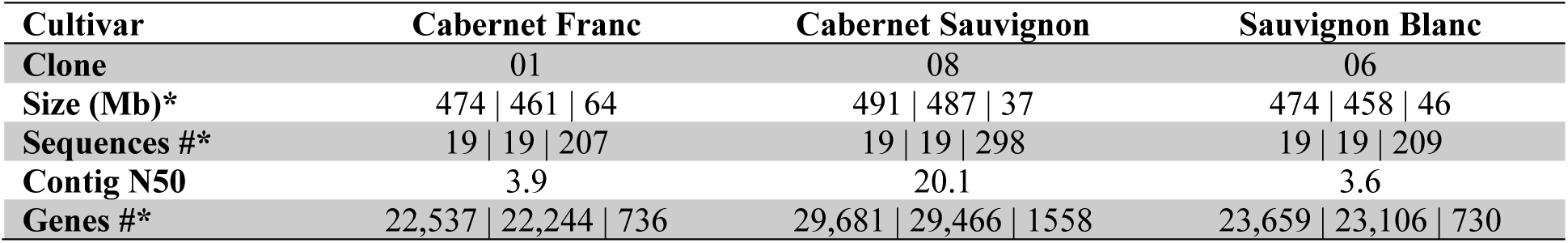
Assembly and annotation statistics for the three reference genomes. *: Values for haplotype 1, 2, and unplaced sequences respectively. For CS, hapCF, hapSB, and unplaced are represented.

We constructed whole-genome bisulfite sequencing (BS-Seq) libraries for CF, CS, and SB, each represented by three individual clones considered here as biological replicates (Table S1). The overall average bisulfite conversion rate was 99% (Table S2) for the three cultivars. The preparation of the genomes and the mapping were performed with bismark [15]. To prevent any reference bias, BS-Seq reads were mapped onto their corresponding reference genome (e.g. CS reads were mapped on the CS assembly). High-quality sites were obtained by filtering out low-confidence cytosine sites (coverage < 10) and merging common sites present in the three clones for each cultivar. Methylation levels were also estimated directly from the HiFi reads for technical validation and showed strong support to the short-read sequencing data independently of the chemistry used (Pearson’s correlation *r* = 0.94, *P* < 2.2^e-16^).

### Distribution of the methylation in the trio of phased genomes

From the BS-seq data analyzed with the corresponding reference per cultivar, an overall range of 20-24.7 Mb total cytosine sites were considered among the different clones accounting for a similar proportion of CG, CHG, and CHH contexts (Table S3). About 73% of the sites were CHH contexts, while CG and CHG only represented 12 and 16% over the 19 chromosomes of CF, CS, and SB (Table S3). Independently of the methylation status, a comparable distribution within and outside the gene space was observed for the three contexts and the three cultivars, with about 80% in the intergenic space, 11.7% in introns, and 8.3% in exons (Figure 1A). Taking into consideration the methylation level, methylated cytosines (presenting at least 20% of methylation) in CG (mCG) and CHG (mCHG) contexts were the most abundant, while CHH showed significantly lower methylation overall (Figure 1B). For all the contexts, the highest percentage of methylated sites was detected in the intronic and intergenic regions. A remarkably high number was observed in the exons for the CG context. Relative to gene loci, a noticeable drop in methylation level was detected in the proximity of the transcription start site (TSS) in the three cultivars (Figure 1C), followed by an increase reaching its highest approximately in the middle of the genes, then decreasing toward the transcription termination site (TTS). Overall, CGs are the most methylated sites in genes followed by CHG. The CHH levels were almost not detectable (Figure 1C).

**Figure 1.**
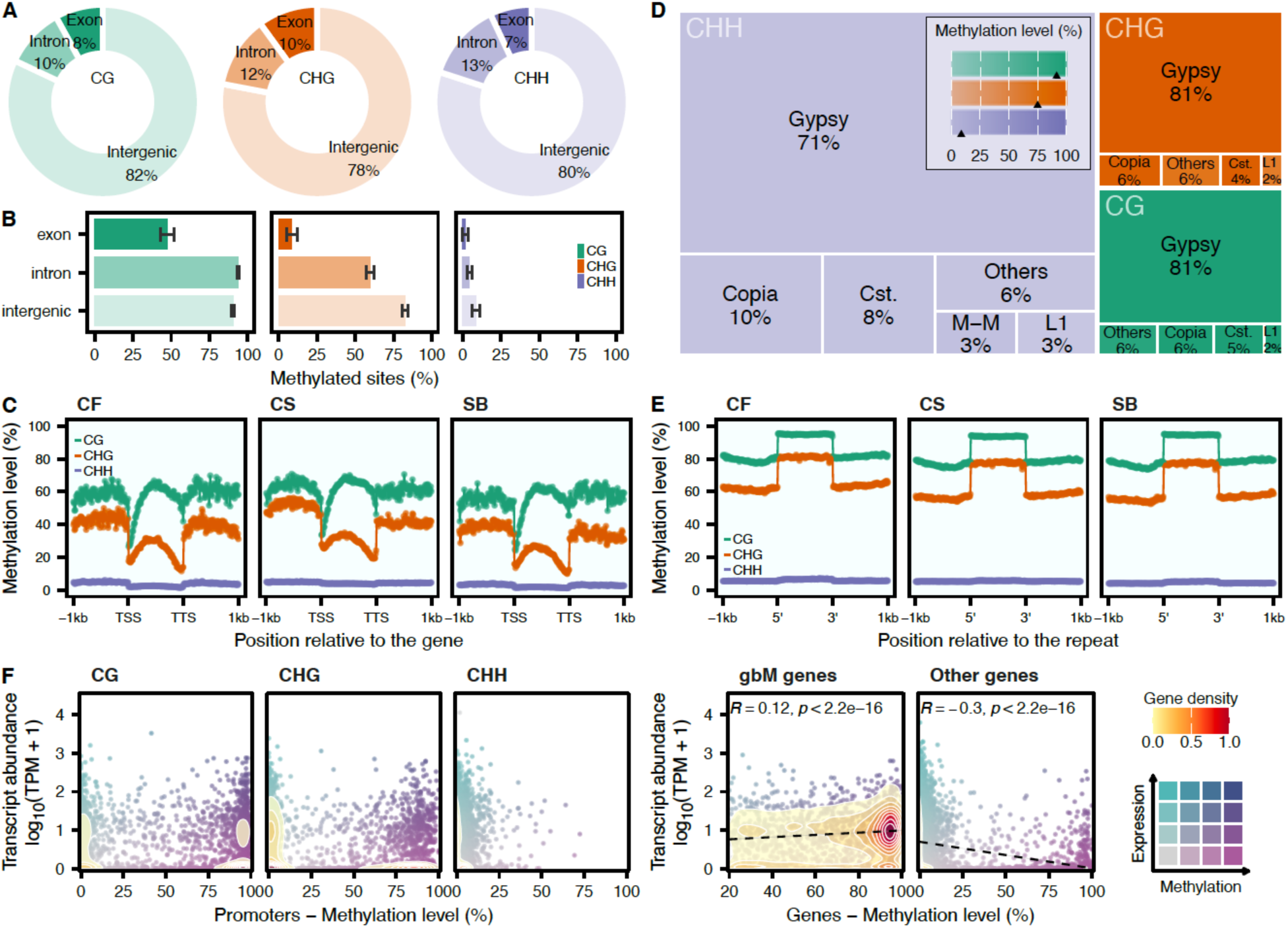
mCGs, the most abundant methylated cytosines, are negatively associated with gene expression except for gbM genes. **A.** Distribution of the total number of methylation contexts relative to the gene space (n = 9 clones). **B.** Proportion of methylated sites (methylation level >=20%) per context group, mean ± sd (n = 9 clones). **C.** Methylation level relative to gene loci. The points represent the average methylation values per bin (n = 3 clones). The -1kb upstream, the gene loci, and the +1 kb downstream sequences were each divided into 100 bins. **D.** Treemap of the total number of methylation contexts in repetitive sequences. The transparency level represents the average methylation level per repeat class. In the legend key, the average methylation level per context is represented by a black arrow. **E.** Methylation level relative to repeat sequences. The points represent the average methylation values per bin (n = 3 clones). The bins were obtained as described for the genes. CG, CHG, and CHH contexts are colored in panels A-E in green, orange, and purple, respectively. TSS: Transcription Start Site; TTS: Transcription Termination Site. Cst.: Custom, as defined by repeatmasker. **F.** Gene expression relative to methylation level in promoter regions and in genes. For genes, Pearson’s correlation coefficient and the associated p-value are represented at the top. gbM: gene body methylation.

In repetitive sequences, CGs, CHGs, and CHHs were found to be distributed similarly with LTR/Gypsy and LTR/Copia being the predominant classes (Figure 1D). Once again, the overall methylation levels were significantly higher in CG (91.4%) and CHG (74.9%), while only 8.6% was detected for the average methylation level in CHH. The levels detected in repeats were relatively higher when compared with the genes (Figure 1E). The shift in methylation relative to the proximity to the repeat sequence was found to be very defined with a noticeable jump in methylation level accounting for a 20% increase in average for CG and CHG sites (Figure 1E). While being the most abundant context in the genome, CHH sites were characterized by a very low methylation level almost unnoticeable in the repeats.

The BS-Seq data were integrated with RNA-Seq data to evaluate the impact of methylation on gene expression. For CS, CG methylation in promoter regions (Figure 1F) was mostly found in genes presenting very low or no expression (40.28% with TPM <1) (Table S4), while for CF and SB, mCG were detected in a similar proportion in genes that were expressed or not (∼30%). For the three cultivars, the lowest proportion of mCHGs was found in expressed genes. For CF and SB, most of the CHGs were not methylated and located in expressed genes. The context CHH was predominantly not methylated for the three genomes. For methylation occurring within gene loci, we observed a more defined impact on gene expression. Genes characterized by gene body methylation (gbM) were mostly expressed while a negative correlation was observed between gene expression and methylation for the other genes overall (Figure 1F, right panel).

### The trio pan-methylome revealed parental conservation of the methylation distribution in Cabernet Sauvignon

Although analyzing the methylation results per genome prevented single-reference genome bias (Figure 2A), performing comparative analyses among cultivars remains challenging. To overcome single-genome limitations, a sequence graph was built to represent each haplotype of the three cultivars with nf-core/pangenome [16,17] (Figure 2B). Within the sequence graph, windows were defined to compare methylation among the embedded paths (haplotypes) corresponding to nodes not longer than 200bp as defined as the optimal window size for the subsequent differential methylation analysis (DMA) (see Materials and Methods section for details). Nodes containing at least one methylation context were then categorized into core (present in all haplotypes), dispensable (shared between two cultivars), and private class (only present in one cultivar).

**Figure 2.**
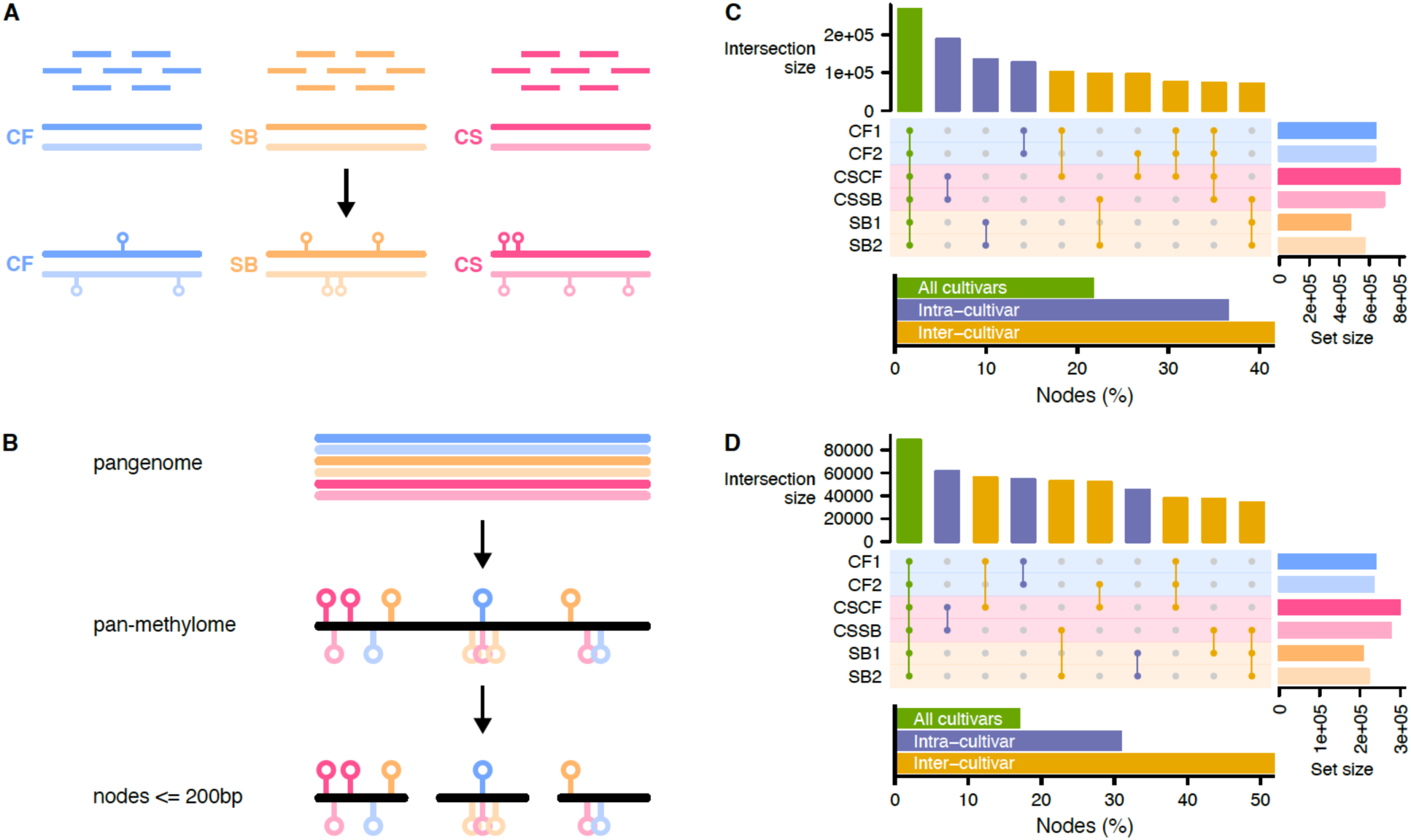
The pan-methylome captured inter-cultivar relatedness. **A.** Cultivar-specific mapping of the bisulfite sequencing data. Each cultivar was represented by 3 individual clones. **B.** Inferring the pan-methylome using a sequence graph. The cultivar-specific diploid methylation data were projected onto the graph. Nodes were chopped to not be longer than 200bp. **C.** Classification of the nodes containing cytosines. The upset plot represents the number of nodes for the top ten most represented combinations of haplotypes. Within these top ten combinations, the nodes proportion per cultivar class (all: node found in all haplotypes; intra: nodes found in the haplotypes of the same cultivar; inter: nodes in haplotypes of more than one cultivar) is represented on the bottom barplot. **D.** Classification of the nodes containing methylated cytosines. The configuration of the plot is the same as in C.

Among the ten largest groups of nodes, the core nodes were the most abundant (Figure 2C), followed by the nodes shared between the haplotypes of the same cultivar (private). Overall, the proportion of nodes detected within the same cultivar and between cultivars was similar. Considering the methylation status, we then defined nodes to be shared between haplotypes only if they presented the same methylation pattern. The methylation-aware distribution of the node groups was found to be enriched for inter-cultivar nodes (Figure 2D). For CF and SB, more nodes were found to be shared with CS than between their own haplotypes (e.g. more methylated nodes between CF1 and CSCF than for CF1 and CF2). While many nodes were shared between haplotypes for CS (i.e. between CSCF and CSSB), the number of nodes shared with the parental haplotypes was proportionally significantly higher when methylation was considered. Altogether, the distribution of methylation patterns between CF and SB, and the phased haplotypes of CS suggest that part of its allele-specific methylome was explained by the methylation found in its parents. It is worth noting that among the combinations of shared nodes from the sequence graph, more than 96% of the shared sequences present also the same methylation status. A good example to confirm the potential inheritance of methylation patterns was to investigate the sex-determining region (SDR) which is known to be highly conserved at the genomic level [18]. Most of the methylated sites were located between the *TREHALOSE-6-PHOSPHAT*E (*T6P*) and the *INAPERTURATE POLLEN 1* (*INP1*) encoding genes (Figure S1), a region known to be very repetitive [19]. By evaluating the methylation patterns, we were able to infer unambiguously the parent of origin for each of the SDR alleles in CS. CSCF (Figure S1B, left panel) was derived from CF hap1 (Figure S1A, left panel) and CSSB (Figure S1B, right panel) was inherited from SB hap2 (Figure S1C, right panel). The analysis of this conserved genomic region clearly supports that the allele-specific methylation observed in the CS progeny was inherited from the parents.

### The differential methylation analysis in regions characterized with allele-specific methylation revealed an overall hyper-methylation pattern shared by CS and CF

We leveraged the structure of the graph to perform differential methylation analysis (DMA) by extracting comparable regions among the embedded paths. By projecting 200bp regions from CS onto the other haplotypes, the corresponding ranges of nodes were extracted and compared (Table S5). Overall, there were more differentially methylated regions (DMRs) when the parent cultivars were compared together (CF vs SB) than with CS (Figure 3A). There was a similar number of hyper-/hypo- DMRs in CG, while in CHG an overall pattern of hypo-methylation was observed in SB. For CHH, regions were mostly hypo-methylated in CS and hyper-methylated in SB when compared to CF (Figure 3A). The distribution of the DMRs within the gene and repeat space tended to follow the general trend (Figure 3B). Most of the DMRs were detected in introns and repeats (32.6% and 35.7% respectively). Very few DMRs were detected for CHH in promoters and exons (7.2% and 1.6% respectively). Among the repeats, the transposable elements (TEs) distribution was different among the comparisons (Figure 3C). CS compared to SB was the only comparison that did not comprise any LTR/Copia in the CG DMRs. Instead, a higher abundance of LINE/L1 (36.4%) was detected for this comparison when compared to the two others. The distribution of TEs for the CHG and CHH DMRs was relatively similar with the LTR/Gypsy being preponderant (∼ 50%). The TIR/CACTA class was only found in the CG DMRs. By comparing the DMR sets, we identified regions separating the parents by DMA and grouped them into two sets named inherited DMR (iDMR; DMR conserved between CS and only one of its parental cultivars): iCF; for conserved DMRs in both CS vs SB and CF vs SB, and iSB; for common DMRs between CS vs CF and SB vs CF (Figure 3D). Hyper-methylated iCF were about three times as much abundant as the hypo-methylated iDMRs (411 hyper- / 141 hypo-methylated). On the other hand, iSB were characterized by a balanced differential (241 / 281 iDMRs). Interestingly, a conserved pattern was observed for both iCF and iSB in promoter regions (Figure 3E), while a reversed pattern characterized the iSB in exons. For introns and repeats, the pattern of the iDMRs demonstrated a clear pattern of allelic-specific methylation for the regions shared between CS and CF when compared to SB.

**Figure 3.**
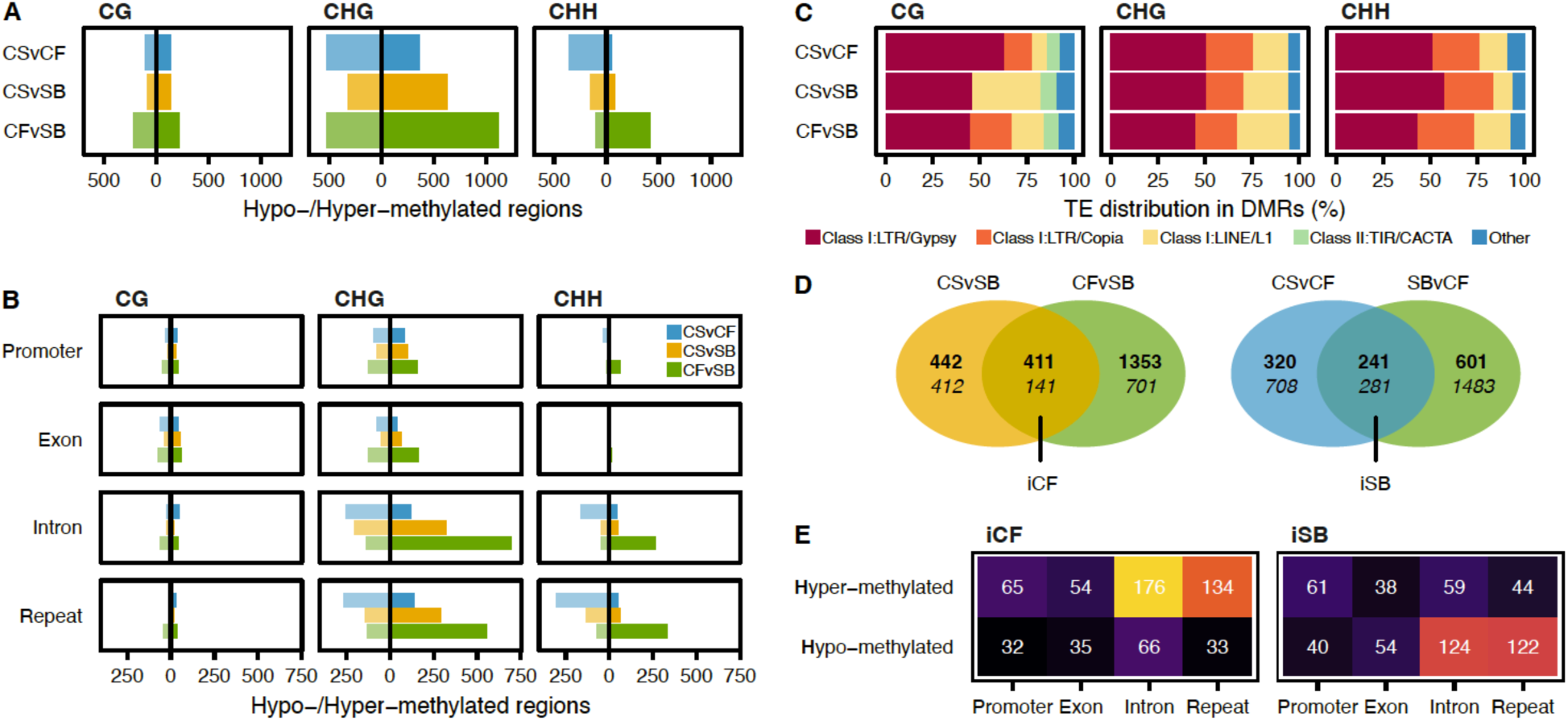
Patterns of methylation from parent lineages are conserved in CS. **A.** Number of hypo-/hyper-methylated regions per methylation context and comparison. **B.** Distribution of the differentially methylated regions (DMRs) per comparison in the gene space (promoter, exon, and intron) and the repeats. **C.** Distribution of the transposable element (TE) content within the DMRs. **D.** Venn diagram representations of the differential methylation analyses and creation of the iCF and iSB subsets. The top numbers are hyper-methylated regions; bottom numbers are hypo-methylated regions. **E.** Distribution of the iCF and iSB DMRs in the genes and repeats.

### Assessing the impact of genomic variants on methylation through linear and graph-based approaches

While allele-specific methylation can strictly be attributed to differences in methylation levels, it can also originate from underlying genomic variants. To evaluate the impact of variants on the DMA, we performed three distinct analyzes: i) a methylation quantification was performed using a single genome as reference to estimate the potential biases of such an approach; ii) the variations embedded in the sequence graph were extracted and their distribution within DMRs was evaluated; iii) the graph was genotyped by mapping short-reads from each sample to assess the clonal variability and the impact of polymorphic sites on methylation differential.

To evaluate the biases inherent to the use of a single-genome reference, we restricted the BS-seq data analysis to the CS genome only. Looking at the mapping of the CF and SB samples, we observed a clear bias towards their corresponding haplotype (e.g. more methylation was detected when CF samples were mapped onto CSCF; Figure 4A). To explore whether underlying genetic variants can explain such pattern, we had to produce an unbiased version of the mappings. To reach this goal, we designed a homology-based correction strategy based on region homology. We defined homologous regions by sequence alignments and ported the corresponding methylation values (i.e. if we use CSCF has a reference, the mappings of SB samples onto CSSB are ported by homology to be represented onto CSCF) (Figure 4B). By comparing the results before and after homology-based corrections, we highlighted a significant recovery of methylation levels (Figure 4C). By further categorizing the polymorphisms between CS haplotypes, we observed a striking enrichment for C>T substitutions inherently biasing the interpretation of DMCs by being undifferentiable from the bisulfite treatment (Figure 4D). Overall, these results unambiguously confirmed that methylation level estimation strongly depends on the correct representation of the genomic diversity for the samples studied and that the use of a linear single reference is not optimal.

**Figure 4.**
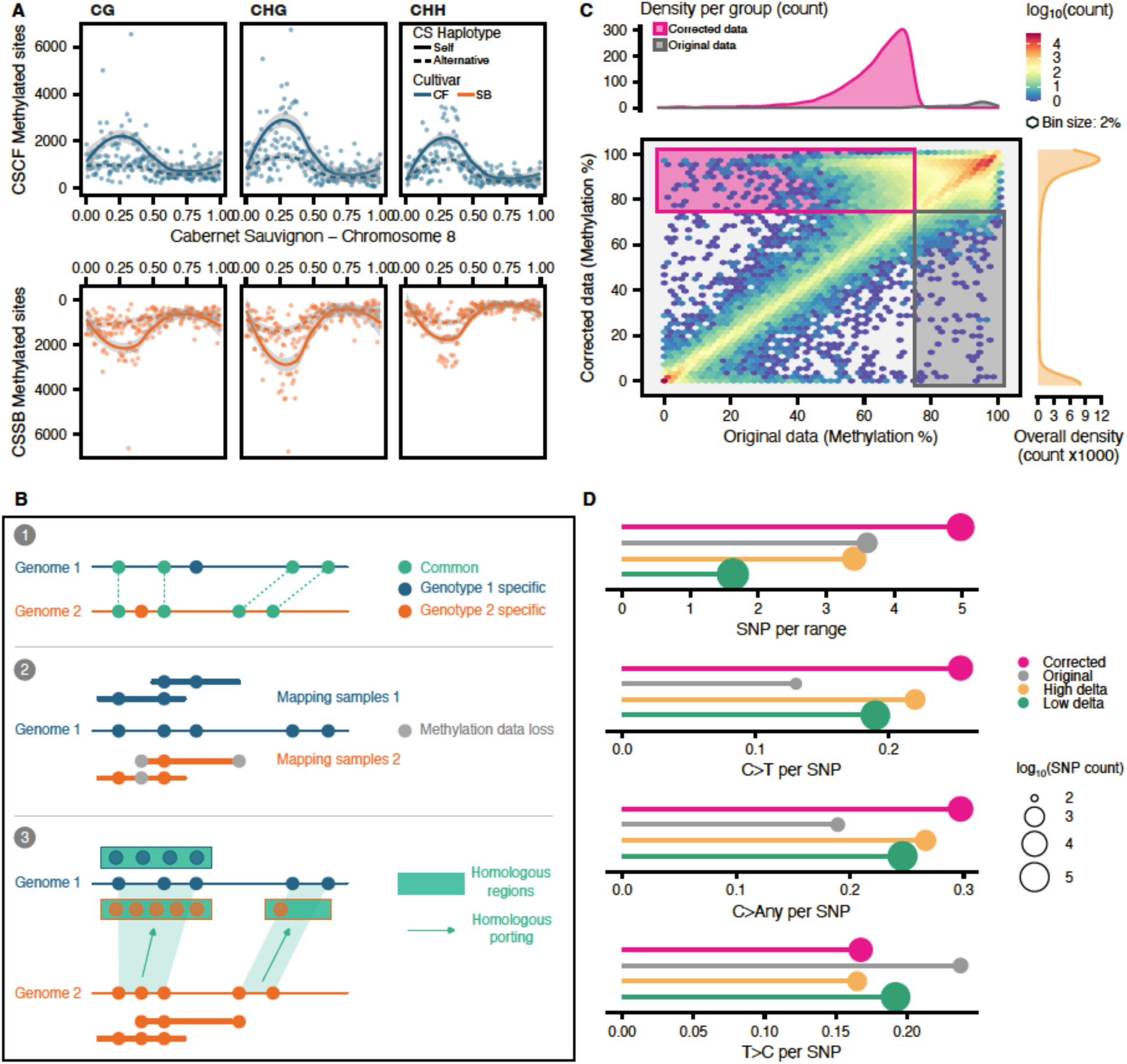
Methylation analysis using a single-genome reference is biased. **A.** Number of methylated sites per context on CSCF haplotype (top panel) and CSSB (bottom panel) for CF (blue) and SB (orange) samples. The chromosome 8 was selected as a representative example. **B.** Correction strategy using homozygous regions: there are genome-specific methylation sites (1) that are not represented when data from multiple genomes are mapped onto a single reference (2). After identifying homozygous regions, methylation estimations can be corrected (3). **C.** Comparison of the methylation levels between the original data and the corrected data. The CG context and the CSCF reference were selected as a representative example. Each axis is divided into 50 bins. The frequency of regions detected within each bin is represented by the filling color gradient. The pink and the grey boxes cover regions where the methylation level is above 75% for the corrected and original data, respectively. The top density plot represents the frequency of regions in these boxes. The right density plot represents the overall density of the regions analyzed. **D.** Distribution of polymorphisms and substitution types on the corrected regions. High deltas represent differences greater than or equal to 20% for the average methylation level between corrected and original data. Low deltas are differences below 20%. Circle size is proportional to the number of SNPs.

From the sequence graph, we extracted the embedded variants to evaluate how they impact the DMA results. For each region compared during the DMA, polymorphic sites for the pairwise comparison were considered (i.e. if the comparison is CF vs SB, a site had to present a different genotype between CF and SB). Sites showing the same genotype were discarded, their presence originated from the graph deconstruction as it outputted variants for all the haplotypes. Variant sites were detected in a higher proportion of regions (84.3%) when the parent cultivars were compared (CF and SB, Table S6), 10% higher on average than the comparisons involving CS (74.6%). For the three methylation contexts, the distribution of variant types was similar with SNPs representing about 60% of the polymorphisms (Figure S2). Among the substitution types, G>A and C>T accounted together for almost 50% of the SNPs (Table S7). No significant differences were observed for the distribution of substitution types within or outside DMRs. In other words, the graph-inferred regions defined for the DMA were not enriched for a specific variant type which appeared to be the main limitation when identifying DMCs with the single-genome approach described above. Variants not categorized as SNPs, INDELs, or MNPs were found to be the second most abundant (Other; 32%, Figure S2). They represent the nested variation structures within the top-level variation bubbles reported by the variant projection from the graph.

Going beyond the three genomes used to build the sequence graph, we considered the genetic variability of each clone by mapping DNA-Seq short reads onto the sequence graph and performed a variant genotyping. After normalizing and filtering the variant sites, we focused on the SNPs as they were the most abundant variant type and retained 3,095,637 sites. For CF and SB cultivars, most of the SNPs where identical among the three clones (Table S8). Although a higher specificity was detected for the clone CS08 as it was the accession used as reference for variant representation, almost 70% of SNPs were common for the CS clones (Table S8). Furthermore, a very similar number of sites were shared when pairwise comparing the clones of the two other cultivars (Table S9), highlighting that despite one clone was selected as reference for the sequence graph construction, it did not impact the mapping nor the variant detection for the others. Overall, SNPs were more stable among clones showing between 10 to 30% more conservation than methylation sites for which an average of 61% of the cytosines were observed in the three clones of each cultivar (Table S3: Normalized/United sets). To understand whether these SNPs had a significant impact on the DMA, we analyzed the distribution of polymorphic sites within the windows that were pairwise compared. No differences were observed when comparing the SNP-containing regions to the overall distribution (Student’s t-test; *P* value > 0.05), the presence/absence of SNPs was not enriched within DMRs (Table S10). Furthermore, among the regions with polymorphic sites, no differences were observed in the number of SNPs for regions considered DMRs or not significant (Table S11). Taken together, these results showed that the sequence graph was able to represent the clonal genomic diversity for each cultivar. The DMRs were not strictly explained by polymorphisms such as the C>T observed with the linear approach and rather represent regions with a true differential of methylation by incorporating genetic variations if there is any.

### Construction of a spliced pangenome graph to integrate transcriptomics with methylation

Methylation within or around gene loci can ultimately impact their expression. To integrate our transcriptomic data with the inferred pan-methylome, a spliced pangenome graph was constructed (Figure 5A). By combining the gene exon-intron junctions of the three different cultivars and the sequence graph, reads can be directly mapped onto the splice pangenome graph which recapitulates both the genomic variants and the transcriptomic splicing information [20](Figure 5B). We observed a great consistency between the genome-level RNA-Seq analysis and the graph-based results (Figure 5C). Moreover, mapping on the sequence graph increased the mapping rate by 19% on average (Table S12) as reads have a larger sequence repertoire to align to. Differential expression analysis (DEA) revealed a higher number of up-regulated genes in CS (about 3 times more) when compared to either CF or SB (Figure 5D). The comparison of CF and SB led to the highest number of differentially expressed genes (DEGs), with a similar number of up- and down- regulated genes (8302/8342 respectively). Interestingly, a large proportion of the up-regulated genes in CS presented the same pattern in the comparison opposing CF and SB (Figure 5E). By comparing the DMA results to the DEA, two noticeable characteristics were observed (Figure 5F): i) DMRs in exons were associated with a higher differential for gene expression overall when compared to DMRs occurring in promoters, ii) for the CHH context, DEGs were detected almost exclusively for DMRs occurring in promoter regions.

**Figure 5.**
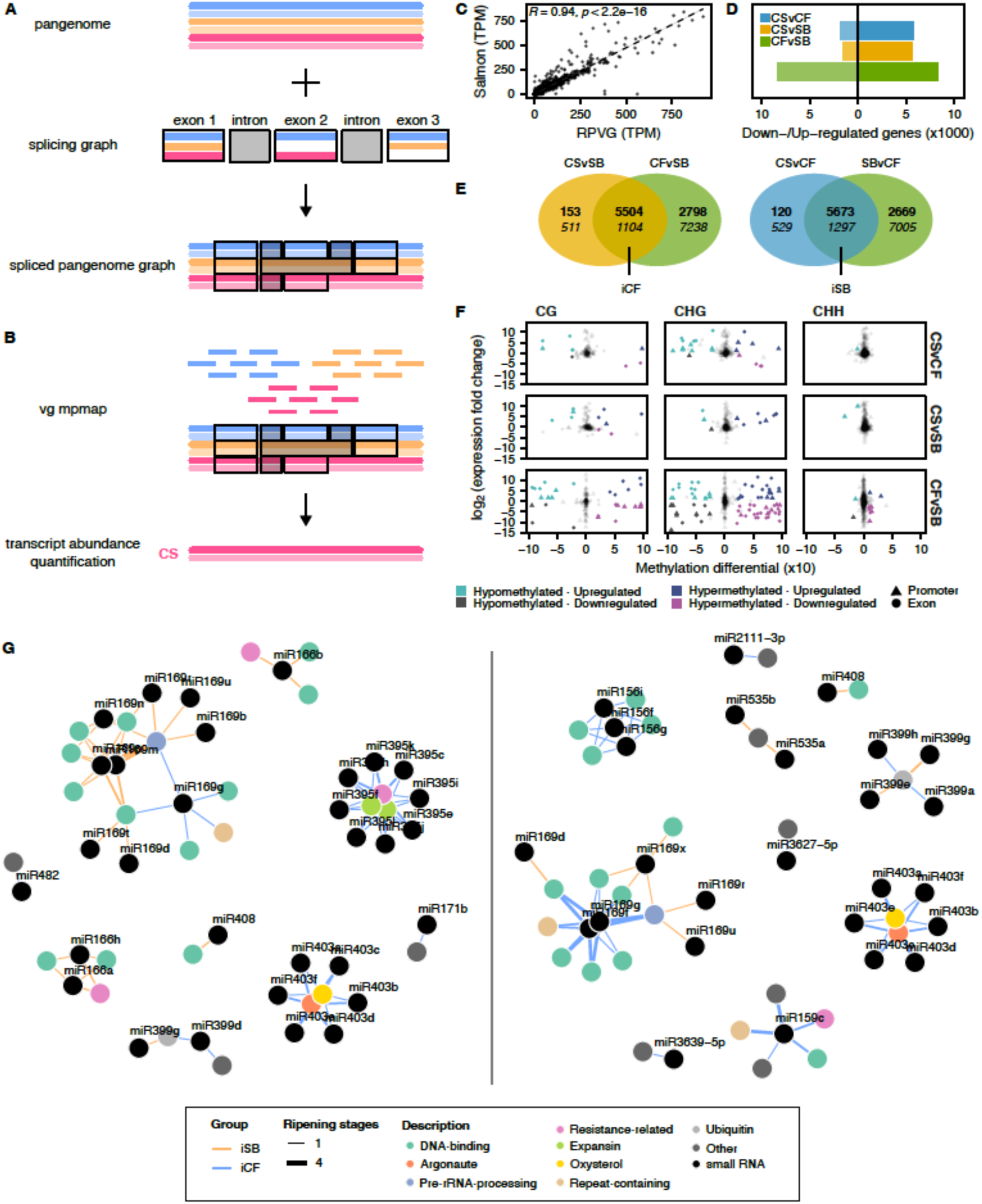
The spliced pangenome graph enables the integration of transcriptomics with methylation and small RNAs. **A.** Creation of a spliced pangenome graph resulting from the combination of the sequence graph and a splicing graph. **B.** Mapping of the RNA-Seq reads directly onto the spliced pangenome graph and transcript abundance quantification against a selected linear reference. **C.** Correlation between transcript abundance quantified using Salmon and RPVG. **D.** Number of differentially expressed genes per comparison. **E.** Venn diagram representations of the differential expression analyses and creation of the iCF and iSB subsets. The top numbers are up-regulated genes, bottom numbers are down-regulated genes. **F.** Comparison of the methylation differential to the expression fold changes per methylation context and comparison. Four groups with different colors were defined based on their differential values for methylation and expression. Triangles represent promoters, and circles represent exons. **G.** miRNA-mRNA network for miRNA precursors (left) and mature miRNAs (right). Colored vertices (genes) and black vertices (miRNAs) are connected by colored edges (iSB: orange; iCF: blue). The edge thickness represents the frequency at which the pairs were detected among the different ripening stages.

To further explore the potential conservation of transcriptomics signatures inherited from CF or SB in CS, we integrated our graph-based results with small RNA (sRNA) sequencing of the same leaf samples. We observed a typical distribution of the small interfering (si)RNAs with the 21-nt located within exons while the 24-nt were found in repetitive regions and promoters (Figure S3). The number of micro (mi)RNAs clusters detected with this approach was very limited (92). Among the roles attributed to siRNAs, their cis activity has been previously documented to regulate methylation [21]. By comparing the location of the 3,605 annotated siRNAs in CS with the previously identified DMRs, only 23 were found to overlap. Among them, 10 showed significant differential expression conserved between CS and CF or SB (Table S13), but this pattern was not concomitantly observed for the methylation.

Since this first approach led to a very limited exploration of the miRNAs class, the pipeline nf-core/smallrnaseq [22] and the database miRbase 22.1 [23] were used to quantify miRNA precursors and mature miRNAs. We also expanded the experimental design to include berry samples at different ripening stages for the three cultivars. After performing DEA for both genes (Figure S4A) and miRNAs (Figure S4B-C), we compared the sets of DEGs and differentially expressed miRNAs (DEmiRs) to identify iDEGs and iDEmiRs (i.e. genes or miRNAs that have a common expression differential between CS and only one of the two parental genotypes). On average, 76% of the DEmiRs had a least one of their targeted genes concurrently differentially expressed. The number of iDEGs and iDEmiRs was then compared and classed by differential (Figure S4D-E). Interestingly, for iCF most miRNA-mRNA pairs showed a negative relationship (i.e. if the miRNA is down-regulated, the targeted gene was found up-regulated) while for iSB the proportion of positive and negative pairs was more balanced (Figure S4F). By building a miRNA-mRNA network for iCF and iSB negative pairs, for the miRNA precursors (Figure 5G, left panel) and the mature miRNAs (Figure 5G, right panel), we observed inherited patterns. Most of the targeted genes were involved in DNA binding functions, mostly transcription factors. Some miRNA precursors were only detected for iCF with for example different forms of miR395 and miR403 targeting the same genes involved in biotic stress resistance, argonaute, and expansin related functions. The miR169 network was the most abundant in targeted iDEGs for both iCF and iSB, however, for mature miRNAs, the network was significantly more stable across berry ripening stages in CF and CS (iCF). Also, the miR403 miRNA-mRNA network was detected exclusively for iCF in the mature miRNAs. Altogether, this network analysis revealed that numerous miRNA-mRNA negative pairs co-exist in CS while they were exclusively detected in CF or SB.

## Discussion

This study demonstrates that despite centuries of clonal propagation, epigenetic marks can be conserved to a certain extent, long after the original parental inheritance. Since cultivars undergoing many cycles of vegetative propagation can accumulate somaclonal epimutations [24,25], each of the cultivars was represented here by three distinct clones.

Methylation is known to be one of the most sensitive omics to the choice of the reference [6]. Single-reference approaches introduce several layers of bias, from impeding the read mapping to limiting the analysis to reference-specific methylation sites and gene annotations. Such a strategy ultimately impairs the proper differential methylation analysis and makes it nearly impossible to distinguish allele-specific patterns. Comparing methylation regions between distinct genomes was challenging, but the correct phasing of the assemblies and the use of the graphical format overcame these limitations. We have not only demonstrated that regions can be defined within a sequence graph to compare methylation across haplotypes (Figure 2B) but also confirmed that the use of a single-reference genome led to strongly biased results (Figure 4) as reported in previous work using a coordinate mapping between divergent genotypes [26]. The sequence graph generated in this study represented the clonal variations for each cultivar by genotyping the embedded polymorphisms. However, simplifying the complexity of such a graph could provide a better representativeness of the clonal variability by incorporating the new polymorphisms detected after mapping but it is computationally challenging [7]. Our results also showed that by corroborating results from short-read data, the direct detection of methylation from long-read sequencing allows the acquisition of high-quality methylomes in parallel of whole genome sequencing with no additional cost [27].

A higher number of genes methylated in CG and CHG contexts showing low or no expression was observed for the cultivar CS (Table S4), averaging around 10% more than CF or SB relative to their total gene contents. Hybrids often show non-additive methylation patterns – progeny’s methylome is not just an average of the parents. Methylation can be instead reinforced, especially in regulatory regions, suggesting epigenetic dominance or methylation gain in hybrids [28,29]. Such a pattern was independently supported by the sequence graph with CS having the highest number of private methylated nodes (Figure 2D). Interestingly, while a high similarity was expected for the intra-cultivar nodes (shared between the haplotypes of the same cultivar genome, Figure 2C), the shared methylated regions were more numerous between the corresponding parent-progeny haplotypes displaying an enrichment for the inter-cultivar nodes when methylation status was considered. Independently of sequence differences, these methylation patterns supported that a parental signature was conserved within the progeny methylome.

To further explore to what extent we could detect allele-specific methylation, we performed many differential analyses to compare the parental lineages and the hybrid (iCF and iSB, Figures 3D and 5E). A very stable result was observed throughout our analyses. The most variable methylation context, over-represented in the iCF and iSB sets, was CHG and mostly occurred in repetitive regions, particularly in TEs (Figure 3). Non-CG methylation is commonly found near transposable elements [1] making it a good candidate for varietal epigenetic differences. In our comparisons, such a signature was characterized by a hyper-methylation of the CHG iDMRs for CS and CF when compared with SB (Figure 3E). The methylation status of CGs was more stable among the genomes (Figure 3A and 3B) however, a few DMRs in TIR/CATCA DNA transposons were specifically detected for this context (Figure 3C). Furthermore, CG DMR overlapping *Copia* LTR retrotransposons only occurred when CF was involved in the comparison. While involved in a significantly lower number of DMRs, these results do not exclude the important contribution of the mCGs in allele-specific methylation. On the other side, methylation was almost undetectable in the CHH context for the three cultivars (Figure 1). The configuration process of the DMA parameters reflected this pattern (Figure S5) with an optimal threshold set twice lower (10%) than for the other methylation contexts (20%). The overall low CHH methylation at the genome level was observed previously in grapes and other plant species [3,30], and is generally expected in clonally propagated species [5].

Overall, our sequence graph strategy helped us to mitigate the biases of reference-based methods. We further expanded its usage by integrating splicing information and performed a pantranscriptomic analysis [20] (Figure 5A and 5B). The stronger methylation differential observed for CHG (Figure 3) was associated with a higher number of DEGs when gene loci had CHG DMRs in their exons or promoter regions (Figure 5F). By investigating the groups of DEGs- DMRs (Figure 5F), we observed that their relationship was not strictly correlated and more nuanced than when we compared expression levels directly to methylation levels (Figure 1F). The correlation between methylation and expression is known to be influenced by many factors including the precise position of the methylation marks within the genic regions [31] (Figure 1C), but also the proximity to repetitive sequences notably transposable elements (Figure 1E) [32]. Furthermore, CS transcriptome was found to express more genes (at higher levels) than CF and SB (Figure 5D). The majority of the DEGs observed when CF and SB were compared were also shared with CS suggesting that the allele showing higher expression was retained. Such a pattern of allele-specific expression was observed previously in other plant models and leads to non-additive gene expression patterns within hybrids [33]. The pantranscriptome was further integrated with small RNA-Seq data (Figure 5G). In addition to DNA methylation, post-transcriptional regulation by sRNAs can indeed play a critical role in modulating gene expression. Our analysis of miRNA–mRNA expression pairs revealed that the sequence graph was able to capture that CS inherited distinct post-transcriptional regulation patterns from each parent. These parent-specific miRNA interactions may have contributed to the differential expression of target genes in CS and reflect regulatory divergence beyond direct transcriptional or epigenetic control.

In conclusion, we demonstrated here that a sequence graph provides a powerful framework for studying methylation inheritance in clonally propagated species like grapes and is relevant for species far beyond the *Vitis* genus. By integrating genomic variation from phased genomes, the sequence graph allowed for the representation of both common and rare alleles, including structural variants and splicing information, both crucial for multi-omics integration to understand epigenetic regulation. Such an approach revealed the parental contributions to epigenetic modifications in their progeny providing insights into the stability and heritability of methylation marks in a model hybridized about 300 years ago.

## Materials and Methods

### DNA sequencing

HiFi sequencing libraries for CS08 were prepared following the manufacturer’s protocol for the SMRTbell Express Template Prep Kit 3.0 (Pacific Biosciences, CA, USA). Libraries were size-selected using the BluePippin system (Sage Science, Beverly, MA, USA) with a cut-off range of 10–50 Kb. After size selection, the HiFi libraries were cleaned up using 1X (v/v) AMPure PB beads (Pacific Biosciences, CA, USA). The concentration and final size distribution of the libraries were evaluated using the Qubit 1X dsDNA HS Assay Kit (Thermo Fisher Scientific, Waltham, MA, USA) and the Femto Pulse System (Agilent, Santa Clara, CA, USA), respectively. The HiFi library was sequenced on a Revio sequencer at the DNA Technology Core Facility, University of California, Davis, CA, USA. For the HiFi libraries of CF01 and SB06, they were prepared as described previously [34] and sequenced using the Sequel II system. For the additional sequencing of CS clones (CS06, CS08, and CS47) a newer library preparation method was used in combination with the PacBio Revio SPRQ Chemistry. Before shearing, the ∼ 3 μg of gDNA for each clone was run through Short Read Eliminator cleanup on the Hamilton Microlab Prep system following manufacturers manufacturer-recommended automation protocol. This was followed by shearing with the Megarupter 3 DNA shearing cartridge (Diagenode: Cat. No. E07010003). Samples were sheared at 30ng/ul with Speed 29 to achieve an insert size distribution ranging from 17kb to 19kb across the clonal gDNA samples. Following shearing, SMRTbell® libraries were constructed for each sample and size selected using the dilute AMPure PB beads as previously described [35]. Samples were simultaneously sequenced with three independent runs in parallel on the Revio Sequencing System with the SPRQ polymerase (PacBio, PN:103-520-100) and SPRQ Sequencing Chemistry (PacBio, PN:103504-900). HiFi reads with methylation calls were generated on the instrument before downstream tertiary analysis. For the short-read sequencing, libraries and sequencing were performed as described previously [19].

### High-density gene map construction

The CDSs from genes annotated on PN40024 [36] were mapped separately on the haplotypes of selected genomes from grapegenomics.com [37]. Assemblies were selected based on their high contiguity from PacBio HiFi sequencing. The mapping was performed with three tools: GMAP v2019-09-12 [38], BLAT v36x2 [39], and minimap2 v2.17-r941 [40]. Thresholds of coverage and identity maximizing the number of uniquely mapping genes were identified separately for each mapper. For GMAP and minimap, the highest number of uniquely mapping genes were obtained setting a threshold of at least 80% identity and coverage, while for BLAT the minimum threshold had to be increased to 97%. The datasets were intersected to obtain a list of 20,165 gene models common to all methods. The alignment coordinates were compared across the selected haplotypes and the CDSs with consistent order and location across all assemblies were retained. The final gene map consists of 19,049 markers based on the synteny of uniquely mapping gene models.

### Genome assembly and annotation

The genome assemblies of Cabernet Sauvignon clone 08 (CS08), Cabernet Franc clone 01 (CF01), and Sauvignon Blanc clone 06 (SB06) were generated from HiFi reads (Yield: 73.21Gb, 14.04Gb, and 12.27Gb respectively) as described previously [34], by combining the hifiasm outputs [41] with the high-density gene map produced in the present study using HaploSync [12]. The phasing of CS08 was further processed with the DNA sequencing data from the parental lineages to generate the haplotypes CF (hapCF) and SB (hapSB). The genome annotation of CS08 was performed as described in previous work [19]. For Cabernet Franc and Sauvignon Blanc, published annotations of Cabernet Franc clone 04 (CF04)[12] and Sauvignon Blanc clone 01 (SB01)[14] were lifted onto the newly assembled clones CF01 and SB06 respectively using lifton [42] with the parameters “-a 0.95 -s 0.95”. The resulting lifted genes were filtered to only retain models coding proteins presenting 100% identity with the source models using diamond [43] blastp and “--id 100 --query-cover 100 --subject-cover 100”.

### Total RNA and Small RNA sequencing

Total RNA isolation was done using a Cetyltrimethyl Ammonium Bromide (CTAB)-based protocol as described previously [44]. Small RNA was purified from total RNA using the quick RNA miniprep kit (Zymo Research, Irvine, USA) as per the manufacturer’s protocol. RNA and Small RNA concentration and purity were assessed with Qubit (Life Technologies, Carlsbad, CA, USA) and a NanoDrop 2000c Spectrophotometer (Thermo Fisher Scientific, Waltham, MA, USA), respectively. RNA sequencing was performed as described previously [19]. Briefly, library preparation was performed with the Illumina TruSeq RNA sample preparation kit v.2 (Illumina, CA, USA) following the low-throughput protocol. After evaluation of library quantity and quality, the sequencing was performed using an Illumina HiSeq 4000 (DNA Technology Core Facility, University of California, Davis, CA, USA) to produce 100bp-long and 150bp-long paired-end reads for the leaves. For the berries, 100bp-long single-end and paired-end reads were generated. Small RNA libraries were prepared using the Illumina TruSeq small RNA library prep kit (Illumina, CA, USA). A hundred nanograms of purified small RNA was used as starting material and 3’ and 5’ RNA adapters were ligated before the reverse transcription. During the PCR amplification of cDNA constructs, individual barcodes were added (with a total of 16 cycles). The libraries were then purified with 1.6X KAPA pure beads (Roche diagnostics) and size selected with the BluePippin instrument (Sage Science, Beverly, MA, USA) using a 3% agarose gel cassette with a cut-off range of 120-170 bp. Size-selected libraries were cleaned with 1X KAPA pure beads (Roche diagnostics, Mannheim, Germany). Libraries were quantified with Qubit (Life Technologies, Carlsbad, CA, USA) and validated with the Agilent High sensitive kit (Agilent Technologies, CA, USA), then pooled at equimolar bases. Two rounds of sequencing were performed, 150bp paired-end by IDSeq on the HiSeqX, and 100bp single-end sequenced on the HiSeq4000 (DNA Technology Core Facility, University of California, Davis).

### Whole Genome Bisulfite Sequencing (WGBS)

The EZ DNA Methylation Gold kit (Zymo Research, Irvine, USA) was used for sodium bisulfite conversion of DNA before library preparation, each DNA sample was spiked in with 0.5 % unmethylated lambda DNA as internal control. All WGBS libraries were prepared from 100 ng of HMW genomic DNA using a TruSeq DNA Methylation kit (Illumina, CA, USA) according to the manufacturer’s protocol. Paired-end sequencing (2 x 150) was performed on a HiSeq4000 (DNA Technology Core Facility, University of California, Davis).

### Bisulfite sequencing data analysis

Pre- and post-trimming quality controls were performed using FastQC v0.11.8 [45] and summarized with MultiQC v1.10.1 [46]. Samples with 150-bp-long reads were trimmed down to 100 bp using trim_galore v0.6.10 [47] and the parameter “--hardtrim5 100”. Trimmed reads were further filtered for quality with the same tool using the parameters “--quality 20 --length 80”. Read depth was normalized across samples by down-sampling the read libraries to the lowest observed value (82462299, Table S1) using seqtk v1.3-r106 with the flags “sample -s100”. After mapping the reads onto lambda genome using Bismark [15] and parameters “--bowtie2 -N 1”, a bisulfite conversion >= 98% was observed for all the samples (Table S2).

When the BS-seq reads were mapped onto the phased genome of their corresponding cultivar, e.g. CS reads mapped onto the CS genome (Table S14), the parameters “--ambiguous --ambig_bam” were added. Any reads giving an ambiguous mapping and absent from the resulting alignment files were further mapped per haplotype. The low rate of residual ambiguous mapping after this second round supports that those reads mostly originated from homozygous regions (Table S15). The alignments on the phased genomes were separated per haplotype to merge them with the corresponding ambiguous read alignments. Another set of alignments was generated by mapping all the samples on the CS genome only to assess single-reference bias.

Deduplication was performed using deduplicate_bismark from the bismark tool suite (Table S16). Methylation statistics were extracted from the deduplicated files using bismark_methylation_extractor. After assessing methylation bias, the final methylation extraction was performed using “bismark_methylation_extractor --paired-end --gzip --multicore 24 -- buffer_size 12G --CX_context --cytosine_report --comprehensive --bedGraph --ignore 15 -- ignore_r2 15 --ignore_3prime 2 --ignore_3prime_r2 2 --output $extractor_outdir --genome_folder $genome_outdir $bam_output” (Table S17).

### Methylation analysis at the genome level

Methylation data were processed using the R package methylkit v1.29.1 [48] in RStudio running R v4.3.3 [49]. Files were opened using methRead with “mincov = 0”, filtered using filterByCoverage and “lo.count=10”, normalized with normalizeCoverage, and concatenated into a single file using unite and the parameter “destrand=FALSE” (Table S3). After converting the unite objects and the structural annotations of each genome into Granges [50], the relative position of the methylation sites was determined using the function findOverlaps. The regions surrounding gene loci and repeats were binned to average the methylation levels. The upstream 1kb, the gene/repeat loci, and the downstream 1kb were each divided into 100 bins.

### Evaluation of the single-reference genome bias

Since more cytosines were detected by mapping the samples (CF or SB) onto their corresponding CS haplotype (hapCF or hapSB respectively, Figure 4A), the strategy was to define homologous regions so that methylation values could be ported. Only one haplotype can be considered by methylkit for the analysis. For example, if the haplotype hapCF is used, more cytosines will be detected for the CF samples (Figure 4B). To correct this bias, results from the mapping of the SB samples on hapSB were ported from pre-defined homologous regions (Figure 4B). Homologous regions were identified by comparing the CS haplotypes hapCF and hapSB using nucmer from mummer v4.0.0rc1 [51]. Each macro-region matching between both haplotypes was then cut into 200bp chunks and mapped a second time using minimap2 [40]. After discarding any ambiguous mapping, mappings of the CF samples on hapSB and mappings of SB samples on hapCF were corrected. Polymorphisms in homologous regions were extracted using the function delta2vcf from mummer (Figure 4D).

### Optimization of the parameters for the differential methylation analysis

Pairwise comparisons were designed using the function “reorganize” from methylkit. Windows were defined using tileMethylCounts with “win.size” and “step.size” being 100-bp increments from 100 to 2000bp, and “cov.bases” ranging from 1 to 10 (Figure S5). The differential methylation analysis (DMA) was performed for each comparison using calculateDiffMeth and ‘overdispersion="MN"’. Significant DMRs were defined using a qvalue < 0.01 and tested for an absolute differential >= 5, 10, 15, 20, and 25 (Figure S5). DMRs were also identified using DSS [52]. The function DMLtest was used with smoothing=TRUE and the significant DMRs were extracted via callDMR using parameters with fixed values “p.threshold=1e-5, minCG=3”, variable delta values ranging from 0.05 to 0.25 by 0.05, and variable region sizes ranging from 50 to 1000bp (Figure S5). DMRs identified using methylkit were tested for overlaps by using the function reduce from GenomicRanges [50].

### Transcript abundance quantification at the genome level

Reads were trimmed based on quality using trimgalore [47] and the following parameters “trim_galore --quality 20 --length 80 --fastqc --gzip --fastqc_args \"-o fastqc/filtered\" –paired $read1 $read2 --output_dir seq/filtered” (Table S18). Salmon [53] was used to index (kmer size = 31) and quantify transcript abundance. For the three cultivars, each set of reads was mapped onto the corresponding diploid genome. Transcript abundance was summarized at the gene level using tximport [54]. The differential expression analysis was performed using DESeq2 [55].

### Sequence graph construction and variant genotyping

A sequence graph was constructed using the nf-core/pangenome pipeline [16,17] which mostly relies on PGGB [56]. The six haplotypes were considered as distinct genomes (2 haplotypes x 3 cultivars) to construct 19 chromosome-level pangenomes with the following configuration: ‘"wfmash_map_pct_id": 90, "wfmash_segment_length": 10000, "wfmash_n_mappings": 5’. Before mapping on the graph, DNA-Seq short reads were trimmed to not be longer than 100bp using cutadapt v.1.18 [57] and “--length 100 --pair-filter both”. Three high-coverage samples (CF04, CS08, and SB01) were down-sampled to the average coverage of the six other samples (53.15M of reads) using seqtk as described for the BS-seq data. The graph was converted to a vcf using vg deconstruct using CS haplotypes as the reference and a new graph was constructed back using vg construct “-m 1024”. The xg index and the snarls were computed using vg index and vg snarls respectively. After pruning the graph with vg prune and the parameter ‘-r’ a gcsa index was constructed with vg index. The paired-end reads were mapped with vg map, the read support was extracted using vg pack, and finally the genotyping was performed with vg call and the option “- a”. After merging the resulting vcf of each sample with bcftools merge v.1.21 [58], the multi-sample vcf was filtered with bcftools filter “-e ’MAF <= 0.25 || QUAL < 30 || INFO/DP < 10 || INFO/DP > 500’”. The vcf records were then normalized with bcftools norm “--atom-overlaps ’*’ -m-any” and SNPs were extracted with bcftools view “-v snps”. A second round of bcftools filter was performed as described above before the final site annotation with bcftools annotate. Any duplicated sites due to the conversion of MNVs into SNVs were discarded. The final filtered vcf contained 3,095,637 SNPs.

### Pan-methylome inference and pan-transcriptome analysis

Before categorizing nodes, the sequence graph was chopped into nodes not longer than 200bp with odgi chop [59]. To perform differential methylation analysis among cultivars, 200bp windows were defined in the phased genome of CS (CSCF and CSSB). Odgi position [59] was then used to determine the corresponding regions in the other genomes. A spliced pangenome graph was generated using vg [20]. The pggb sequence graph was converted to PackedGraph format with vg convert and chopped to contain nodes not longer than 32bp using vg mod. A non-conflicting id space between the 19 pangenomes was obtained with vg ids before merging them all together using vg combine. The spliced pangenome graph was finally generated using vg rna. Prior to mapping RNA-Seq reads the gcsa and dist indexes were generated using vg index. The mapping was performed using vg mpmap using CS as reference for gene expression. Counts were extracted from the multipath alignments using rpvg [20]. The differential expression analysis was performed following the genome-level pipeline described above.

### Small RNA analysis

Small RNA sequencing data were aligned and annotated using Shortstack v.4.1.1 [60]. The annotated clusters were filtered to only retain valid DicerCalls. To further characterize the miRNA, the nf-core/smallrnaseq pipeline [22] and miRbase 22.1 [23] were used. Counts were processed with DESeq2 [55] for differential expression analysis as described for the classic RNA-Seq analysis.

## Data availability

All the raw sequences were deposited on NCBI SRA under the BioProject number PRJNA1244791. The new genome assemblies and annotations are available on Zenodo (CS, 10.5281/zenodo.15191822; CF, 10.5281/zenodo.15191856, and SB, 10.5281/zenodo.15191869) and hosted on grapegenomics.com [37] with blast and jbrowse2 services. The matches from the LiftOn results between the previous and the new annotations were uploaded in the corresponding repositories (10.5281/zenodo.15191856 and 10.5281/zenodo.15191869). The high-density gene map was uploaded in the Zenodo repository 10.5281/zenodo.15191822. The code and scripts used for the analyses are available on GitHub (https://github.com/noecochetel/Epigenomics_Trio).

## Authors’ contributions

NC, AV, and DC designed the study. RFB processed the samples, performed the nucleic acids extractions, and prepared the sequencing libraries. NC performed all the bioinformatic analyses. JL and PP prepared and sequenced the HiFi libraries. NC and DC wrote the original manuscript draft, and the co-authors reviewed and edited the manuscript. All authors read and approved the final manuscript.

## Acknowledgments

The authors acknowledge the DNA Technology Core at the UC Davis Genome Center for sequencing support; Andrea Minio (Institute for Biomedicine, EURAC Research, Bolzano, Italy) for bioinformatics support; Andrea Guarracino (University of Tennessee Health Science Center, Memphis, TN, USA) for providing access to the *odgi* tool; and Guilherme De Sena Brandine (Pacific Biosciences, Menlo Park, CA, USA) for assistance with methylation extraction from HiFi reads. This work was supported by NSF grant #1741627, the E.&J. Gallo Winery, and the Ray Rossi Endowment in Viticulture and Enology.

## Supplemental Files

**Supplemental File 1: Table S1.** Whole-Genome Bisulfite Sequencing read statistics (number of reads). **Table S2.** Bisulfite conversion statistics (%). **Table S3.** Methylkit data processing (number of cytosines). **Table S4.** Methylation vs gene expression. **Table S5.** DMA results (number of significant hypo-/hyper-methylated regions). **Table S6.** Number of windows with variants. **Table S7.** Substitution type distribution. **Table S8.** Number of SNPs shared between clones. **Table S9.** Number of SNPs shared between pairs of clones. **Table S10.** DMRs with polymorphic sites compared to the overall DMRs. **Table S11.** Average number of SNPs in regions consired DMR or not significant. **Table S12.** RNA-Seq mapping rates. **Table S13.** Differentially expressed sRNA clusters overlapping DMRs. **Table S14.** Bismark results - mapping on phased genomes (number of reads). **Table S15.** Bismark results - mapping per haplotype (number of reads). **Table S16.** Bismark deduplication results (number of alignments). **Table S17.** Bismark results. Un-/Methylated numbers represent the cytosine coverage by the reads. **Table S18.** RNA-Seq read filtering.

**Supplemental File 2: Figure S1.** Methylation patterns are inherited at the sex-determining region. **Figure S2.** Distribution of the graph-inferred variants in the DMRs. **Figure S3.** Distribution of the sRNA clusters in the genome of CS. **Figure S4.** Identification of iDEGs and iDEmiRs in berries. **Figure S5.** Benchmark for differentially methylated regions (DMRs).

